# Age and experience affect the biosynthesis and emission of a Drosophila pheromone

**DOI:** 10.1101/2022.01.27.478004

**Authors:** Jérôme Cortot, Jean-Pierre Farine, Matthew Cobb, Claude Everaerts, Jean-François Ferveur

**Affiliations:** Centre des Sciences du Goût et de l’Alimentation, UMR6265 CNRS, UMR1324 INRA, Université de Bourgogne Franche-Comté, 6, Bd Gabriel, 21000 Dijon, France; School of Biological Sciences, University of Manchester, Manchester M13 9PT, U.K.

**Keywords:** *cis*-Vaccenyl acetate, microbiota, social interaction, preimaginal conditioning, *desat1*

## Abstract

The most studied pheromone in Drosophila melanogaster, cis-Vaccenyl Acetate (cVA), is synthesized in the male ejaculatory bulb and transferred to the female during copulation. Combined with other chemicals cVA can modulate fly aggregation, courtship, mating and fighting. It is not detected on the cuticle of isolated males and is only released by males involved in social or sexual interaction. We explored the mechanisms underlying both cVA biosynthesis and emission in males of two wild types and a pheromonal mutant line. The effects of ageing, adult social interaction, and maternally-transmitted cVa and microbes — both associated with the egg chorion— on cVA biosynthesis and emission were measured. While ageing and genotype changed both biosynthesis and emission in similar ways, early developmental exposure to maternally-transmitted cVA and microbes strongly decreased cVA emission but not the biosynthesis of this molecule. This indicates that the release — but not the biosynthesis — of this sex pheromone strongly depends on early developmental context. The mechanism by which the preimaginal effects occur is unknown, but reinforces the significance of development in determining adult physiology and behaviour.

**Summary statement:** We show how the biosynthesis and release of a key Drosophila pheromone is affected by ageing, by early exposure to this pheromone and to microbes, and by social context.

## Introduction

The volatile *Drosophila* pheromone (Z)-11-octadecenyl acetate (*cis*-Vaccenyl Acetate; *c*VA) is probably the most intensively studied pheromone in the animal kingdom. It is synthesized by *Drosophila melanogaster* males (Bontonou and Wicker-Thomas, 2014; Butterworth, 1969; Guiraudie-Capraz et al., 2007; Symonds and Wertheim, 2005) and can be detected both on the male cuticle and in the ejaculatory bulb. It is transferred from males to females during mating and is then introduced into food during egg-laying, and plays a series of behavioural roles, altering responses to other compounds (Bartelt et al., 1985a; Bartelt et al., 1985b; Hedlund et al., 1996; Jaenike et al., 1992; Schaner et al., 1987). These changing effects of *c*VA when presented in a blend can be seen in two ways. When combined with food compounds, cVA induces aggregation in male and female flies of several Drosophila species (Bartelt et al., 1985a; Keesey et al., 2016; Schaner et al., 1987; Wertheim et al., 2002); the neuronal basis of this effect is beginning to be understood (Das et al., 2017). When combined with the *D*.*melanogaster* male cuticular hydrocarbon 7-T, *c*VA inhibits male courtship (Everaerts et al., 2018; Jallon et al., 1981; Laturney and Billeter, 2016), stimulates female mating and modulates the intensity of *D*.*melanogaster* male-male aggressive behaviour (Wang et al., 2011).

Although *c*VA detection and processing has been intensively investigated at the cellular level, in particular sexually dimorphic effects (attraction and repulsion) (Datta et al., 2008; Kurtovic et al., 2007; Lebreton et al., 2015; Ruta et al., 2010), the factors involved in cVA biosynthesis and emission are poorly understood. *c*VA is produced in the ejaculatory bulb (Butterworth, 1969; Guiraudie-Capraz et al., 2007) and its level increases with age (Bartelt et al., 1985a). It is not released by isolated virgin males and can be detected on the male cuticle only after a physical interaction with another fly of either sex (Everaerts et al., 2010). With one exception (Bartelt et al., 1985a), only superficial levels of *c*VA – those found on the cuticle or at the male genital opening have been studied. There has been no investigation of internal levels of *c*VA and therefore of the biosynthesis of this pheromone.

There is another aspect to behavioural responses to *c*VA – they can be affected by experience. If males court recently-mated females, which are rich in *c*VA, their subsequent courtship of virgin females is reduced in intensity (Ejima et al., 2007; Siegel and Hall, 1979). Developmental factors can shape these courtship responses to *c*VA. When the egg is laid by a recently mated female, it is covered in cVA that the male has introduced together with his sperm and other compounds (Billeter and Wolfner, 2018). If this *c*VA is removed from the egg, the males that eventually emerge show a weaker courtship reduction in response to *c*VA (Everaerts et al., 2018). This suggests either that *c*VA directly affects development in the egg, or that early contact with *c*VA as the larva emerges from the egg shapes the adult response. Furthermore, this pre-imaginal effect also seems to be mediated by other maternally-transmitted factors present on the outside of the egg, such as microbes.

In this paper we explored how the link between biosynthesis and emission of *c*VA changes as male flies age. We also studied the effects of early exposure to *c*VA and to microbial factors on *c*VA biosynthesis and emission and looked at the role of *desat1*, a gene that simultaneously affects the production, detection and processing of pheromones in *Drosophila* and is expressed in a variety of tissues, including the male genital tract (Nojima et al., 2019). Our results show that early experience, in particular exposure to *c*VA and microbes on the egg, increased the superficial amount of the pheromone on the surface of the adult insect, making more of the compound available for immediate detection by other flies. This effect was paralleled by a decrease in the internal level of *c*VA, presumably as the compound was trafficked to the outside of the fly.

## Material and Methods

### Flies

*D. melanogaster* were raised on yeast / cornmeal / agar medium and kept at 24±0.5° with 65±5% humidity on a 12 L : 12 D cycle. Unless indicated, flies were transferred every two days to avoid larval competition and to regularly provide abundant progeny for testing. Flies were isolated under light CO_2_ anaesthesia 0–2h after eclosion. We tested two wild-type stocks, Canton-S (Cs) and Dijon2000 (Di2) and the *desat1*^*1573*^ homozygous mutant (*desat1*) which was produced by the insertion of a PGal4 transposable element in the regulatory region of the *desaturase1* gene (Marcillac et al., 2005). Unless specified otherwise males and females were held separately in fresh glass food vials in groups of 5 flies until they were studied.

### <u>cis</u>-Vaccenyl Acetate (cVA)

Flies were frozen for 5 min at -20°C and then individually extracted following two procedures. To characterize superficial *c*VA, individual flies were immersed for 5 min at room temperature using 30μl hexane. To measure global *c*VA, individual flies were immersed at room temperature for 24 hours in 30μl dichloromethane. In both cases, the solvent solutions contained 3.33 ng/μl of C26 (*n*-hexacosane) and 3.33 ng/μl of C30 (*n*-triacontane) used as internal standards. *c*VA amounts were quantified by gas chromatography using a Varian CP3380 gas chromatograph fitted with a flame ionization detector, a CP Sil 5CB column (25 m x 0.25 mm internal diameter; 0.1 μm film thickness; Agilent), and a split–splitless injector (60 ml/min split-flow; valve opening 30 sec after injection) with helium as carrier gas (50 cm/sec at 120°C). The temperature program began at 120°C, ramping at 10°C/min to 140°C, then ramping at 2°C/min to 290°C, and holding for 10 min. We determined *c*VA amounts on groups of 50 eggs using the hexane-extraction procedure for individual flies.

### Egg and fly manipulation

To obtain enough eggs, about 100 three to four days old mass-reared females of the three strains were allowed to lay eggs for 3 hours in an egg-laying device consisting of a 50mm Petri dish filled with 1mL 3% agar striped with fresh yeast to stimulate egg-laying. After this period, flies were removed and eggs collected. To determine the influence of the egg chorion on *c*VA levels, eggs were collected and rinsed five times in fresh deionized and sterile water. They were then dechorionated by immersion for a few minutes in a 3% solution of sodium hypochlorite, followed by three washes with sterile deionized water. About 50 eggs were transferred in each food vial.

For the D1 vs D5 experiment, eggs laid by females who had mated either one day (D1) or five days before (D5) were transferred to plain food in groups of 50.

In the case of preimaginal exposure to *c*VA-rich food, groups of 50 intact eggs were transferred onto regular food with added synthetic *c*VA (15ng/mm^3^; Cayman Chemical, Ann Arbor, MI, USA (50 mg/ml solution in ethanol, purity > 98%);(Everaerts et al., 2018)). Adult males eclosing from this food were immediately transferred onto plain food before being tested.

To determine the effect of bacteria on *c*VA levels, we kept the 3 strains for two generations on food containing an antibiotic mixture (50μg/mL tetracycline + 200μg/mL rifampicin + 100μg/mL streptomycin; (Sharon et al., 2010)).

For all adult experiments, males produced by the various treatments were collected immediately after adult eclosion and kept in groups of 5 flies (unless specified otherwise) until they were subjected to *c*VA extraction. To determine the effect of adult social interaction, males eclosing from the control experiment were kept in groups of 5 flies, or alone, and tested when 4-days old.

### Statistics

All statistical tests were performed using XLSTAT 2021 (Addinsoft, 2021). We compared the amount of *c*VA in the males of the 3 genotypes at all ages using Kruskall-Wallis test followed by Conover-Iman multiple pairwise comparisons (p=0.05, with a Bonferroni correction). The effect of each treatment was estimated by comparison with the corresponding control using the Mann-Whitney test, for each genotype at each age.

## Results

The amount of global *c*VA (this corresponds to the sum of *c*VA on the cuticle and in the ejaculatory bulb) can be considered as a indirect measure of *c*VA biosynthesis. In male flies global *c*VA showed a gradual increase with age, with the main increase being observed in the first day (Figure 1A – note that for the sake of clarity, on all figures the amount of *c*VA is presented on a log scale; statistical analysis was performed on untransformed data). Overall there were no clear and consistent differences between the three genotypes (the two wild-types – Cs and Di2 – and the *desat1* mutants) for this measure, with all three strains reaching around 1000 ng of *c*VA per fly at around 4 days old. These levels were maintained not only in 12 day old flies (Figure 1A) but over subsequent weeks of life, up to 40 days in the case of Cs males (Figure S1). Clear differences were observed for the amount of *c*VA detected via a shorter extraction protocol, which measured the superficial *c*VA present on the cuticle and at the genital opening and therefore available for immediate detection by other flies (Figure 1B). This superficial *c*VA can be considered as the amount of *c*VA emitted by a male, and is evidently dependent on biosynthesis, as measured by the global level of *c*VA. Overall, *desat1* mutant males showed significantly lower levels of superficial *c*VA than the two wildtype strains, indicating that the mutation affects the emission of *c*VA, perhaps by altering hydrocarbons secreted on the inner surface of the genital tract. Cs males emitted the highest amounts at all ages.

**Figure 1.**
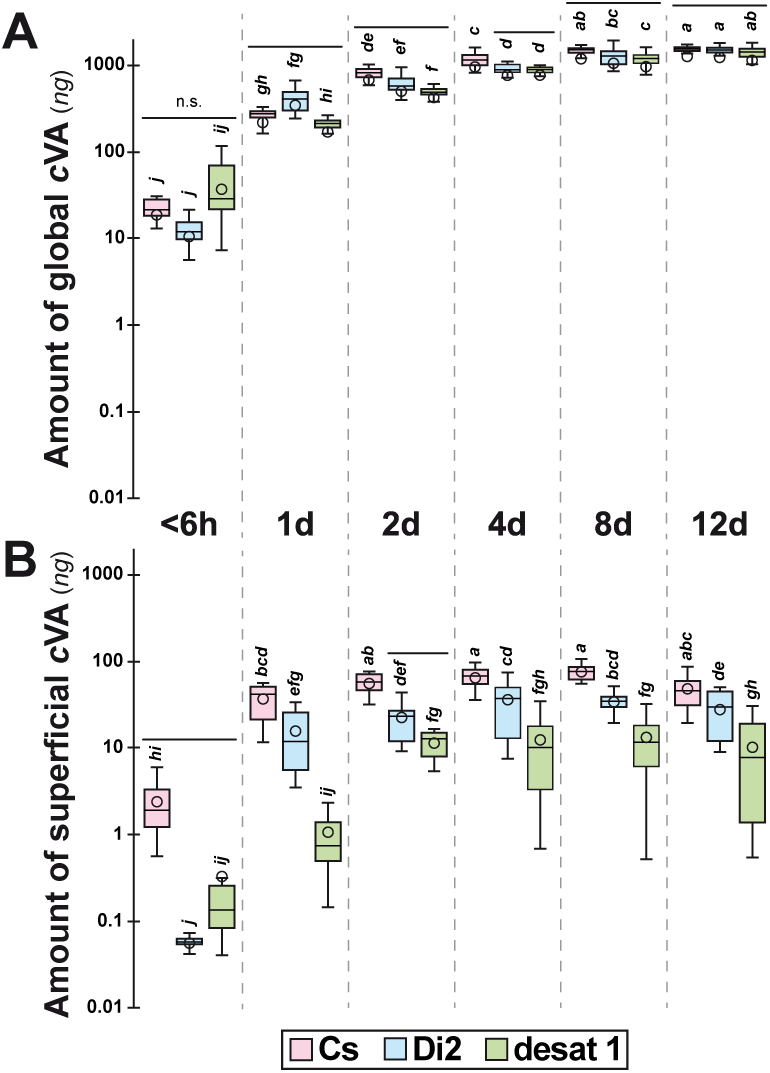
Age–related variation for global and superficial amounts of *cis*-Vaccenyl Acetate in male flies. *cis*-Vaccenyl Acetate (*c*VA) was measured (in ng) in males of the following ages (indicated between the two series of graphs): less than 6 hours (<6h), 1 day (1d), 2 days (2d), 4 days (4d), 8 days (8d) and 12 days (12d). We measured both the global amount of *c*VA (**A**; obtained after a 24-hour extraction with dichloromethane) and the superficial amount of *c*VA (**B**; obtained after a 5-min hexane extraction). We compared these values in males of three genotypes: Canton-S (Cs), Dijon 2000 (Di2) and desat1^1573^ homozygous mutant (desat1). Data are shown as box and whisker diagrams. The lower and upper box edges represent the first and third quartiles, while the median value is indicated by the inner small horizontal bar and the mean by the plain dot. The ends of the whiskers shown below and above each box represent the limits beyond which values were considered anomalous. For each series of graphs, we compared all values including the three genotypes at all tested ages using a Kruskall-Wallis test followed by Conover-Iman multiple pairwise comparisons (p=0.05, with a Bonferroni correction). *N* = 15 for each sample. Horizontal bars indicate no statistical difference between genotypes at each age while different letters indicate significant differences for all genotypes and ages considered.

We next investigated how early experience affected these patterns of *c*VA synthesis and emission. When eggs are laid by a recently-mated female, they are covered with two factors that may alter the behaviour of the adult fly – *c*VA and bacteria. *c*VA is transferred into the female during mating and is introduced into the medium along with the eggs, while the female also has her own microbiome, some or all of which is transferred onto the eggs before or during egg-laying. Removing the chorion from eggs, a process that eliminates both these factors, led to a slight but significant and consistent increase in the amount of global *c*VA produced by the two wild-type strains throughout adult life (Figure 2A). However, from two days old onwards, the amount of superficial *c*VA immediately available on the cuticle and at the genital opening of males showed slight but consistently significantly lower levels in wild-type males from dechorionated eggs (Figure 2B). This suggests that early exposure to chorion-associated factors allows the consistent trafficking of cVA to the outside of the fly; in the absence of this exposure, less *c*VA was emitted. No effect of dechorionation was observed for *desat1* mutants (Figures 2A, 2B), which showed consistently lower levels of superficial *c*VA compared to the wildtype males, but no significant difference for global *c*VA levels.

**Figure 2.**
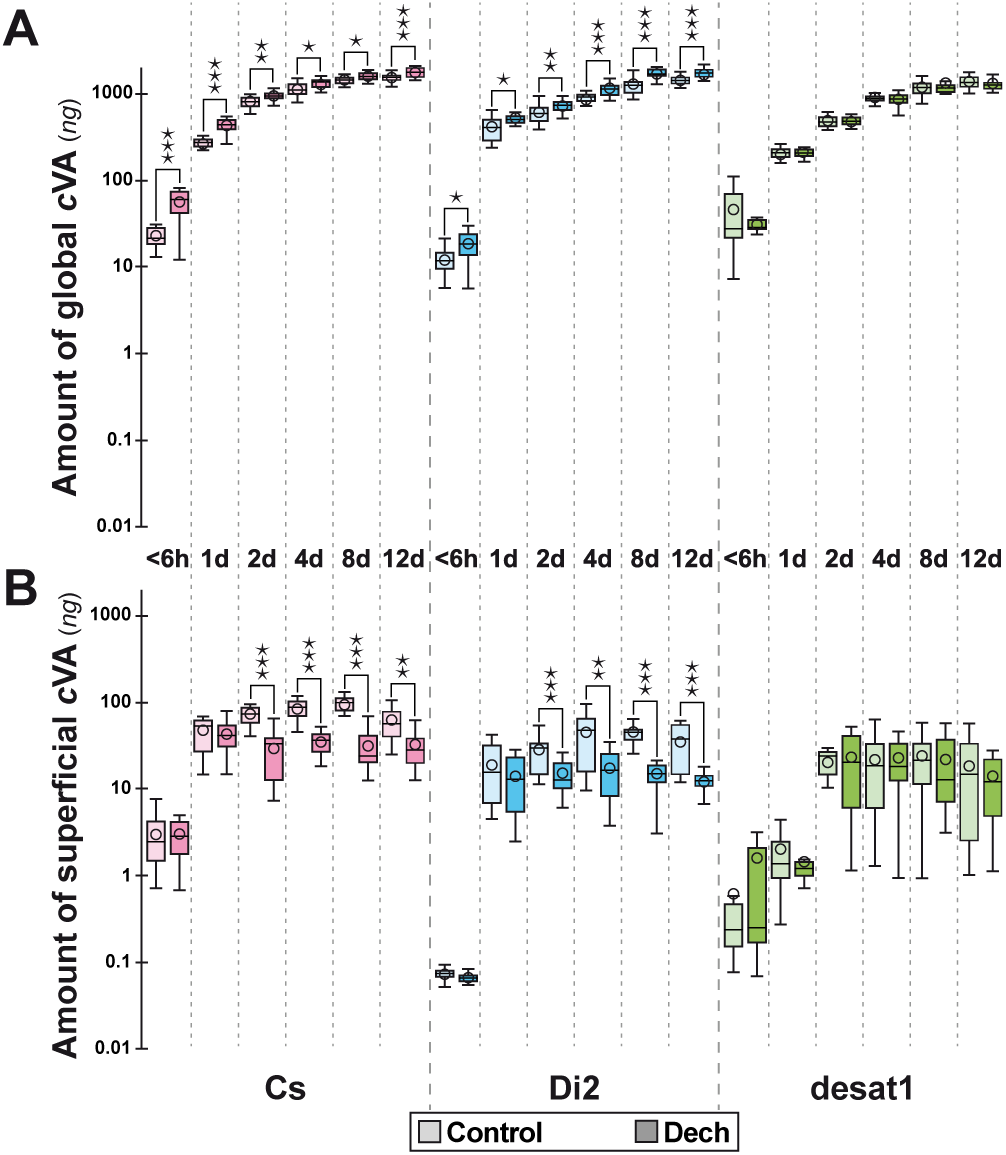
Age-related effect of dechorionation on global and superficial amounts of *c*VA. The amounts of (**A**) global and (**B**) superficial *c*VA were compared between males from dechorionated eggs (“Dech”) and from intact eggs (Control). We compared males of the three genotypes and ages as those described in Figure 1. We used the Mann-Whitney test to compare the dechorionation effect for each genotype at each age: ***: p<0.001; **: p<0.01; *: p<0.05. The absence of stars indicates that no significant difference was detected. *N* = 15 for Cs and desat1 flies; *N* = 13-15 for Di2 flies.

To separate the effect of early exposure to *c*VA from that of exposure to bacteria, we next reared flies on antibiotic-rich food for two generations. The effects were less clear-cut, but there were small age-related variations in the amount of global *c*VA and the amount of superficial *c*VA produced by wild-type males (Figures 3A, 3B) that had been reared on antibiotic-rich food, compared to controls. Some significant differences were observed between the two conditions for *desat1* mutant flies, in particular in the first six hours of life where control levels of *c*VA were significantly higher than those for antibiotic-treated flies. These findings suggest that the effect of early exposure to maternally-transmitted factors on *cVA* levels does not only involve bacteria.

**Figure 3.**
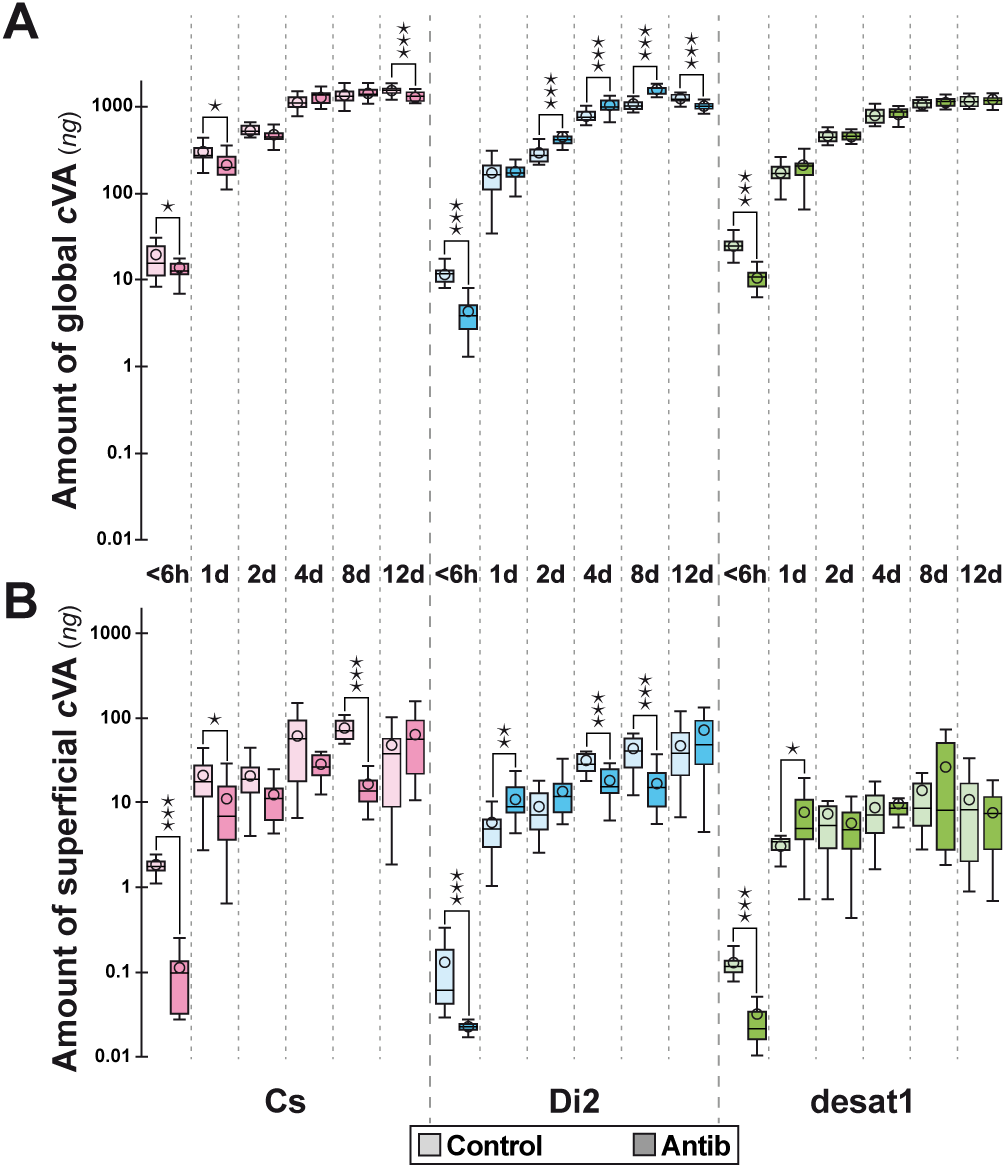
Age-related effect of antibiotics on global and superficial amounts of *c*VA. The amounts of (**A**) global and (**B**) superficial *c*VA were compared between males raised on regular food containing antibiotics (see Material and methods) for two generations. These treated males (“Antib”) were compared with males raised on plain food (Control) of three genotypes and at different ages (see Figure 1 caption). For statistics, see Figure 2 caption. *N* = 15.

To explore the role of *c*VA in this effect, we indirectly manipulated the amount of *c*VA present on the surface of the egg. We first measured the amount of *c*VA present on eggs laid at 1 and 5 days after mating (D1 and D5, respectively). *c*VA levels on the eggs declined significantly between the two ages, for all three genotypes (Figure 4A). By comparing the amounts of *c*VA produced by males that emerged from D1 or D5 eggs, we were therefore able to explore the quantitative impact of early *c*VA exposure on adult male *c*VA production. All three genotypes showed occasional but non-consistent significant differences, unlike the clear effect induced by dechorionation. There were differences between the two wild-type strains: variations of global and superficial *c*VA levels in Cs males were more similar to those induced by dechorionation than in Di2 males (compare Figure 4 with Figure 2). The effects of low exposure to *c*VA in *desat1* males were not consistent but did show some significant differences (Figure 4), whereas no exposure to *c*VA resulted in no differences (Figure 2). These data suggest that exposure to reduced *c*VA level present on the eggs at 5 days after mating was not sufficient to affect the amount of *c*VA emitted by adult males in a consistent manner. Taken together these three experiments suggest that the emission of *c*VA in adult male flies is regulated by pre-imaginal exposure to maternally-transmitted factors – *c*VA and microbes.

**Figure 4.**
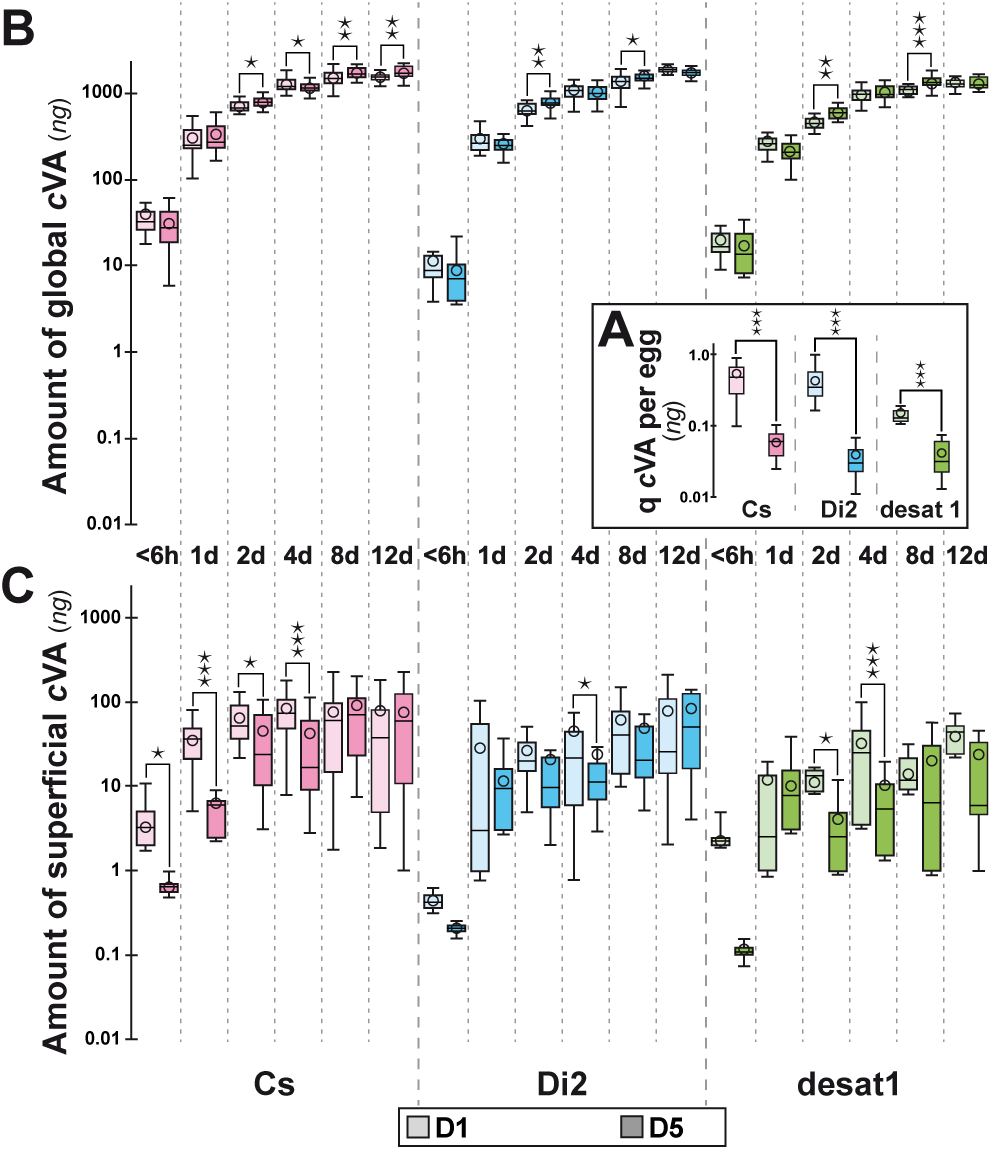
Effect of egg exposure to maternally-transmitted *c*VA. (**A)** The amount of *c*Va was measured in groups of 50 eggs either laid one day (“D1”) or five days (“D5”) after copulation. The amounts of (**B**) global and (**C**) superficial *c*VA were also determined on adult males originating from D1 and D5 eggs. We compared eggs and adult males of the Cs, Di2 and desat1 genotypes (see Figure 1 caption). For statistics, see Figure 2 caption. *N* = 13-15 groups of eggs (A); *N* = 15 for adult samples (B & C).

Finally, the social control of the biosynthesis and emission of cVA was explored by comparing the amount of global and superficial *c*VA on males that were held in groups of five flies, as in the rest of our study, or singly. There were no differences in the biosynthesis of *c*VA, as measured globally (Figure 5A), but in all three strains there were significantly lower levels of superficial *c*VA in males that were reared alone (Figure 5B), highlighting the role of social interactions in the emission of *c*VA.

**Figure 5.**
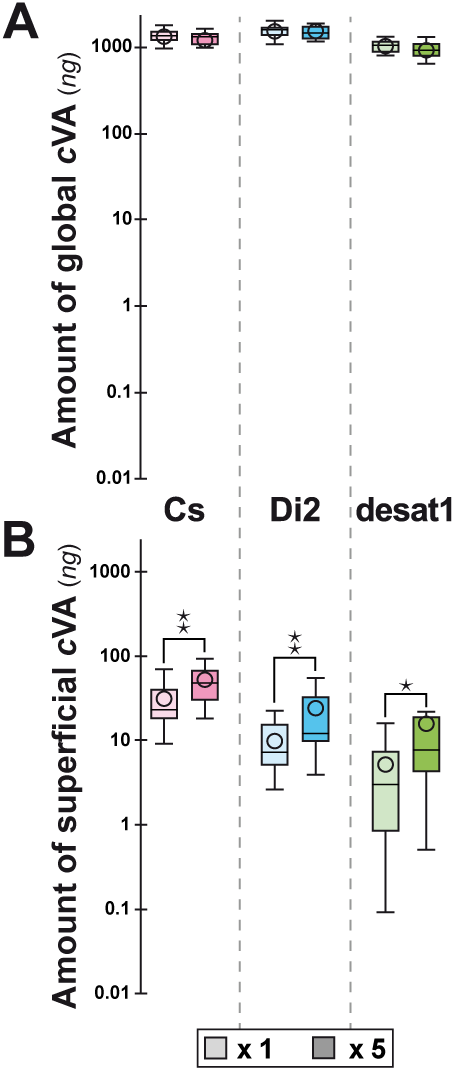
Effect of adult grouping on global and superficial amounts of *c*VA. The amounts of (**A**) global and (**B**) superficial *c*VA were compared between four-days old males kept either isolated (“x1”) or grouped (“x5”) during adult life. For genotypes, ages and statistics, see Figure 1 and Figure 2 captions. *N* = 15.

## Discussion

This study is the first exploration of the production and emission of a *Drosophila* pheromone. We show that while there is an evident necessary relation between these two factors (emission requires biosynthesis), it is not reciprocal – material can be synthesised but not released. This is partly a consequence of our experimental design: males were housed together, allowing for the emission of small quantities of *c*VA during male-male interactions, but had no contact with females, leading to an accumulation of *c*VA inside the males, as none was introduced into females during mating.

We were able to compare production and emission by using extremely different extraction durations – 5 minutes for superficial extraction, which will have measured material on the cuticle and in the genital tract, and 24 hours for measuring global *c*VA levels, which will have included both superficial levels and *c*VA that had been synthesised in the ejaculatory bulb. These two aspects correspond to different functions of *c*VA – global *c*VA measures material that could be transferred to a female during mating, altering the responses of male flies to her and enabling her to attract other flies to the food site where she has laid her eggs, while superficial *c*VA corresponds to material that could immediately be detected by another fly of either sex, playing a role in aggression and courtship. The global amount of *c*VA increased with age, as previously reported by (Bartelt et al., 1985a). The increase in the amount of *c*VA seen between 6 hours and 8 days could be due to the accumulation of *c*VA in the ejaculatory bulb (Guiraudie-Capraz et al., 2007). Conversely, the amount of superficial *c*VA remained more or less constant after 2 days. Superficial levels of *c*VA on the cuticle can be detected by SPME fibres only after mating (Everaerts et al., 2010; Farine et al., 2012).

Studies of other insects may shed light on this balance between synthesis and emission. In many moths, female pheromones are synthesized in an abdominal gland and released with a circadian rhythmicity (for example in *Trichoplusia ni, Pectinophora gossypiella, Argyrotaenia velutinana* ; (Groot et al., 2014; Jurenka, 2017)). Pheromone synthesis depends on the pheromone biosynthesis activating neuropeptide (PBAN; (Jurenka, 2017)), while pheromone emission — usually associated with “calling” behaviour (Allison and Cardé, 2016) — is controlled either directly from the terminal nerve input (Christensen et al., 1994) or indirectly through the muscular contraction of the gland (Raina et al., 2000; Solari et al., 2007). In female cockroaches, secretory cells may synthesise pheromones during periods of sexual receptivity but regress during sexual inactivity, such as gestation (Schaal et al., 2003). A number of neuroendocrine factors such as PBAN, ecdysteroids and juvenile hormone can regulate pheromone synthesis (Blomquist and Vogt, 2003; Rafaeli, 2002; Tillman et al., 1999). Neuronal signals either descending from the central nervous system or ascending through the ventral nerve cord can also modulate sex pheromone synthesis and/or emission (Teal et al., 1989). The situation may be different in Drosophila males which appear to synthezise *c*VA continuously while they emit *c*VA only during encounters with conspecifics (Everaerts et al., 2010; Farine et al., 2012; Laturney and Billeter, 2016; Wang et al., 2011). In contrast, cuticular hydrocarbon biosynthesis depends on the circadian activity of the *desat1* gene (Krupp et al., 2008).

Further investigation of the neuroendocrine control of *c*VA physiology will be necessary, but our results already show that there are significant developmental factors involved. This can be seen through our observation that the levels of *c*VA are affected by very early experience. Removing the chorion from eggs, thereby eliminating both the paternal *c*VA that was present in the female’s genital tract and had covered the eggs as they were laid, and maternal microbiota, led to a slight but significant and consistent increase in the amount of global *c*VA produced by males. This was shown by both wild-type strains throughout the life of the fly. We were able to explore the relative roles of paternal *c*VA and maternal microbiome in producing this effect by rearing flies on antibiotic-rich food, and by observing the effects of reducing the amount of *c*VA present on the eggs. Both manipulations altered the global levels of *c*VA, but to a much greater extent than when the eggs were covered with both *c*VA and the maternal microbiome. Because the levels of *c*VA on eggs laid five days after mating were still apparently above the threshold to affect *c*VA emission in adult males, future experiments will be needed to identify this threshold, by reduce *c*VA levels on eggs still further, perhaps by extending gap between mating and laying to 10 days.

Adding synthetic *c*VA to the laying medium produced no effect on either global or superficial *c*Va of resultant wild type males (Figure S2). This suggests that the effect of *c*VA depends on it being maternally-transmitted on the chorion and/or non-homogenously present in the food –the higher levels of *c*VA present on the chorion compared to our experimental treatment may account for the lack of effect we observed. We conclude that the emission of *c*VA in adult male flies is regulated by pre-imaginal exposure to a range of maternally-transmitted factors, including *c*VA and microbes (there may also be other factors that we have yet to identify, which are also removed by dechorionation).

The mode of action of these maternal factors on the developing egg is unknown, but may involve either altering gene regulation and/or acting as modulators of protein function. Further investigation will be required to elucidate this. Our observations are not the only effects of early exposure to *c*VA on adult behaviour: pre-adult exposure to maternally-transmitted *c*VA alters how adult male courtship is affected by *c*VA (Everaerts et al., 2018).

The three genotypes we studied did not show the same responses. The differences between the two wildtype strains – Cs and Di2 – were relatively minor and presumably reflect differences in genetic background. For example, the different responses of the two wildtypes to antibiotic treatment suggests that the two wild types males have a different sensitivity to early exposure to *c*VA and microbes. The differential response of desat1 flies was of far greater biological significance. desat1 flies have the same genetic background as Cs, allowing us to impute differences in their responses to the mutation in desat1. While all males of all three strains showed very close global *c*VA levels, the two wild type males showed higher superficial levels of *c*VA, suggesting that the desat1 mutant interferes with the trafficking of *c*VA from the ejaculatory bulb to the exterior, thereby reducing the emission of the pheromone. This effect could be due to the mutation leading to the absence of desaturated hydrocarbons in the male genital tract, where they may play a role akin to lubrication. The lack of any effect of dechorionation on *c*VA emission in *desat1* males is at least partly because the levels of superficial cVA were already extremely low.

We have shown that the biosynthesis and the emission of *c*VA show different changes with aging and with social context. Furthermore, simultaneous exposure to microbes and *c*VA during egg development, and perhaps during larval life, is involved in the regulation of *c*VA emission in adult males. Our next challenge will aim to investigate the biological mechanisms underlying this effect and to explore the effect of early exposure to *c*VA on female behaviour.

## SUPPLEMENTAL FIGURE LEGENDS

**Supplemental Figure 1. Age-related variation in global amounts of *cis*-Vaccenyl Acetate**. The variation of the global amount of *cis*-Vaccenyl Acetate (*c*VA) was measured (in ng) in males of different ages until 40 days. We compared males of three genotypes: Canton-S (Cs), Dijon 2000 (Di2) and desat1^1573^ homozygous mutant (desat1; see Figure 1). For the sake of clarity, we only show the mean value for each age and genotype. *N* = 5 - 10..

**Supplemental Figure 2. Effects of preimaginal exposure to synthetic *cis*-Vaccenyl Acetate (*c*VA) on global *c*VA**. We compared *c*VA levels in wild type Canton-S (Cs) and Dijon 2000 (Di2) males at various ages. These males underwent preimaginal development either on plain food (“Control”) or on food with added 15ng *c*VA /mm^3^ food (“cVA”). Immediately after adult eclosion, all males were transferred onto regular food. Data are shown as box and whisker diagrams. The lower and upper box edges represent the first and third quartiles, while the median value is indicated by the inner small horizontal bar and the mean by the plain dot. The ends of the whiskers shown below and above each box represent the limits beyond which values were considered anomalous. We used the Mann-Whitney test to compare the food effect at each age and for each genotype. *N* = 15 except for 2- and 4-day old Di2 flies (*N* = 19-45). No effect of preimaginal diet was observed. For more information, refer to the legends of Figures 1 and 2.

## REFERENCES

Addinsoft. (2021). XLSTAT 2021: Data Analysis and Statistical Solution for Microsoft Excel. Paris, France: Addinsoft.

Allison, J. D. and Cardé, R. T. (2016). Variation in moth pheromone: causes and consequences. In Pheromone communication in moths: evolution, behavior and application., pp. 25–41. Oakland, California: University of California Press.

Bartelt, R. J., Jackson, L. L. and Schaner, A. M. (1985a). Ester Components Of Aggregation Pheromone Of Drosophila-Virilis (Diptera, Drosophilidae). Journal of Chemical Ecology 11, 1197–1208.

Bartelt, R. J., Schaner, A. M. and Jackson, L. L. (1985b). Cis-Vaccenyl acetate as an aggregation pheromone In Drosophila melanogaster. Journal of Chemical Ecology 11, 1747–1756.

Billeter, J. C. and Wolfner, M. F. (2018). Chemical cues that guide female reproduction in Drosophila melanogaster. Journal of Chemical Ecology 44, 750–769.

Blomquist, G. J. and Vogt, R. G. (2003). Biosynthesis and detection of pheromones and plant volatiles—introduction and overview. In Insect Pheromone Biosynthesis and Molecular Biology: The Biosynthesis and Detection of Pheromones and Plant Volatiles, eds. G. J. Blomquist and R. G. Vogt), pp. 3–18. Amsterdam: Elsevier Academic Press.

Bontonou, G. and Wicker-Thomas, C. (2014). Sexual communication in the Drosophila genus. Insects 5, 439–458.

Butterworth, F. M. (1969). Lipids of Drosophila: a newly detected lipid in the male. Science 163, 1356–7.

Christensen, T. A., Lashbrook, J. M. and Hildebrand, J. G. (1994). Neural activation of the sex-pheromone gland in the moth Manduca sexta: real-time measurement of pheromone release. Physiological Entomology 19, 265–270.

Das, S., Trona, F., Khallaf, M. A., Schuh, E., Knaden, M., Hansson, B. S. and Sachse, S. (2017). Electrical synapses mediate synergism between pheromone and food odors in Drosophila melanogaster. Proceedings of the National Academy of Sciences of the United States of America 114, E9962–E9971.

Datta, S. R., Vasconcelos, M. L., Ruta, V., Luo, S., Wong, A., Demir, E., Flores, J., Balonze, K., Dickson, B. J. and Axel, R. (2008). The Drosophila pheromone cVA activates a sexually dimorphic neural circuit. Nature 452, 473–477.

Ejima, A., Smith, B. P. C., Lucas, C., Van Naters, W. V., Miller, C. J., Carlson, J. R., Levine, J. D. and Griffith, L. C. (2007). Generalization of courtship learning in Drosophila is mediated by cis-vaccenyl acetate. Current Biology 17, 599–605.

Everaerts, C., Cazalé-Debat, L., Louis, A., Pereira, E., Farine, J. P., Cobb, M. and Ferveur, J. F. (2018). Pre-imaginal conditioning alters adult sex pheromone response in Drosophila. PeerJ eCollection 2018.

Everaerts, C., Farine, J.-P., Cobb, M. and Ferveur, J.-F. (2010). Drosophila Cuticular Hydrocarbons Revisited: Mating Status Alters Cuticular Profiles. PLoS One 5, e9607.

Farine, J.-P., Ferveur, J.-F. and Everaerts, C. (2012). Volatile Drosophila Cuticular Pheromones Are Affected by Social but Not Sexual Experience. PLoS One 7, e40396.

Groot, A. T., Schöfl, G., Inglis, O., Donnerhacke, S., Classen, A., Schmalz, A., Santangelo, R. G., Emerson, J., Gould, F., Schal, C. et al. (2014). Within-population variability in a moth sex pheromone blend: genetic basis and behavioural consequences. Proceedings. Biological sciences / The Royal Society 281: 20133054.

Guiraudie-Capraz, G., Pho, D. B. and Jallon, J. M. (2007). Role of the ejaculatory bulb in biosynthesis of the male pheromone cis-vaccenyl acetate in Drosophila melanogaster. Integrative Zoology 2, 89–99.

Hedlund, K., Bartelt, R. J., Dicke, M. and Vet, L. E. M. (1996). Aggregation pheromones of Drosophila immigrans, D. phalerata, and D. subobscura. Journal of Chemical Ecology 22, 1835–1844.

Jaenike, J., Bartell, R. J., Huberty, A. F., Thibault, S. and Libler, J. S. (1992). Aggregations in mycophagous Drosophila (Diptera: Drosophilidae): candidate pheromones and field responses. Annals of the Entomological Society of America 85, 696–704.

Jallon, J. M., Antony, C. and Benamar, O. (1981). An anti-aphrodisiac produced by Drosophila melanogaster males and transferred to females during copulation. Comptes Rendus de l’Academie des Sciences Serie III Sciences de la Vie 292, 1147–1149.

Jurenka, R. (2017). Regulation of pheromone biosynthesis in moths. Current Opinion in Insect Science 24, 29–35.

Keesey, I. W., Koerte, S., Retzke, T., Haverkamp, A., Hansson, B. S. and Knaden, M. (2016). Adult frass provides a pheromone signature for Drosophila feeding and aggregation. Journal of Chemical Ecology 42, 739–747.

Krupp, J. J., Kent, C., Billeter, J. C., Azanchi, R., So, A. K. C., Schonfeld, J. A., Smith, B. P., Lucas, C. and Levine, J. D. (2008). Social experience modifies pheromone expression and mating behavior in male Drosophila melanogaster. Current Biology 18, 1373–1383.

Kurtovic, A., Widmer, A. and Dickson, B. J. (2007). A single class of olfactory neurons mediates behavioural responses to a Drosophila sex pheromone. Nature 446, 542–546.

Laturney, M. and Billeter, J. C. (2016). Drosophila melanogaster females restore their attractiveness after mating by removing male anti-aphrodisiac pheromones. Nature Communication 7, 12322

Lebreton, S., Trona, F., Borrero-Echeverry, F., Bilz, F., Grabe, V., Becher, P. G., Carlsson, M. A., Nässel, D. R., Hansson, B. S., Sachse, S. et al. (2015). Feeding regulates sex pheromone attraction and courtship in Drosophila females. Scientific Reports 5, 13132.

Marcillac, F., Bousquet, F., Alabouvette, J., Savarit, F. and Ferveur, J. F. (2005). A mutation with major effects on Drosophila melanogaster sex pheromones. Genetics 171, 1617–1628.

Nojima, T., Chauvel, I., Houot, B., Bousquet, F., Farine, J. P., Everaerts, C., Yamamoto, D. and Ferveur, J. F. (2019). The desaturase1 gene affects reproduction before, during and after copulation in Drosophila melanogaster. Journal of Neurogenetics. 33, 96–116.

Rafaeli, A. (2002). Neuroendocrine control of pheromone biosynthesis in moths. International Review of Cytology 213, 49–91.

Raina, A. K., Wergin, W. P., Murphy, C. A. and Erbe, E. F. (2000). Structural organization of the sex pheromone gland in Helicoverpa zea in relation to pheromone production and release. Arthropod Structure & Development 29, 343–353.

Ruta, V., Datta, S. R., Vasconcelos, M. L., Freeland, J., Looger, L. L. and Axel, R. (2010). A dimorphic pheromone circuit in Drosophila from sensory input to descending output. Nature 468, 686–90.

Schaal, B., Coureaud, G., Langlois, D., Giniès, C., Sémon, E. and Perrier, G. (2003). Chemical and behavioural characterization of the rabbit mammary pheromone. Nature 6944, 68–72.

Schaner, A. M., Bartell, R. J. and Jackson, L. L. (1987). (z)-ll-Octadecenyl acetate, an aggregation pheromone in Drosophila simulans. Journal of Chemical Ecology 13, 1777–1786.

Sharon, G., Segal, D., Ringo, J. M., Hefetz, A., Zilber-Rosenberg, I. and Rosenberg, E. (2010). Commensal bacteria play a role in mating preference of Drosophila melanogaster. Proceedings of the National Academy of Sciences of the United States of America 107, 20051–20056.

Siegel, R. W. and Hall, J. C. (1979). Conditioned responses in courtship behavior of normal and mutant Drosophila. Proceedings of the National Academy of Sciences of the United States of America 76, 3430–3434.

Solari, P., Crnjar, R., Spiga, S., Sollai, G., Loy, F., Masala, C. and Liscia, A. (2007). Release mechanism of sex pheromone in the female gypsy moth Lymantria dispar: a morpho-functional approach. Journal of Comparative Physiology A: Neuroethology, Sensory, Neural, and Behavioral Physiology 193, 775–785.

Symonds, M. R. and Wertheim, B. (2005). The mode of evolution of aggregation pheromones in Drosophila species. Journal of Evolutionary Biology 18, 1253–63.

Teal, P. E. A., Tumlinson, J. H. and Oberlander, H. (1989). Neural regulation of sex pheromone biosynthesis in Heliothls moths. Proceedings of the National Academy of Sciences of the United States of America 86, 2488–2492.

Tillman, J. A., Seybold, S. J., Jurenka, R. A. and Blomquist, B. J. (1999). Insect pheromones — an overview of biosynthesis and endocrine regulation. Insect Biochemistry and Molecular Biology 29, 481–514.

Wang, L., Han, X., Mehren, J., Hiroi, M., Billeter, J. C., Miyamoto, T., Amrein, H., Levine, J. D. and Anderson, D. J. (2011). Hierarchical chemosensory regulation of male-male social interactions in Drosophila. Nature Neuroscience 14, 757–62.

Wertheim, B., Dicke, M. and Vet, L. E. M. (2002). Behavioural plasticity in support of a benefit for aggregation pheromone use in Drosophila melanogaster. Entomologia Experimentalis et Applicata 103, 61–71.

